# miRge 2.0: An updated tool to comprehensively analyze microRNA sequencing data

**DOI:** 10.1101/250779

**Authors:** Yin Lu, Alexander S. Baras, Marc K. Halushka

**Author notes:** Corresponding Author: Marc K. Halushka MD, PhD, Department of Pathology, Johns Hopkins University SOM, 720 Rutland Avenue / Ross Bldg. Rm 632B, Baltimore, MD, USA, 21205.

## Abstract

miRNAs play important roles in the regulation of gene expression. The rapidly developing field of microRNA sequencing (miRNA-seq; small RNA-seq) needs comprehensive bioinformatics tools to analyze these large datasets. We present the second iteration of miRge, miRge 2.0, with multiple enhancements. miRge 2.0 adds new functionality including novel miRNA detection, A-to-I editing analysis, better output files, and improved alignment to miRNAs. Our novel miRNA detection method is the first to use both miRNA hairpin sequence structure and composition of isomiRs resulting in a more specific capture of potential miRNAs. Using known miRNA data, our support vector machine (SVM) model predicted miRNAs with an average Matthews correlation coefficient (MCC) of 0.939 over 32 human cell datasets and outperformed miRDeep2 and miRAnalyzer regarding phylogenetic conservation. The A-to-I editing analysis implementation strongly correlated with a reference dataset’s prior analysis with adjusted R^2^ = 0.96. miRge 2.0 comes with alignment libraries to both miRBase v21 and MirGeneDB for 6 species: human, mouse, rat, fruit fly, nematode and zebrafish; and has a tool to create custom libraries. With the redevelopment of the tool in Python, it is now incorporated into bcbio-nextgen and implementable through Bioconda. miRge 2.0 is freely available at: https://github.com/mhalushka/miRge.

## Introduction

MicroRNAs (miRNAs) are short, single-stranded RNAs that post-transcriptionally regulate gene expression via mRNA decay and/or translational repression (1,2). MiRNAs are transcribed by RNA polymerases II and III, generating precursors that undergo a series of cleavage events to form mature miRNAs (3). Around 30% to 60% of all human protein coding genes are regulated by miRNAs (4), involved in almost all biological process ranging from development to metabolism to cancer (5-7).

With the continued popularity of small RNA sequencing to characterize miRNAs, much attention has been focused on miRNA alignment software. In 2015 we introduced miRge, a fast, multiplexing method to align miRNAs and other RNA species to expressed libraries (8). Since that time, a number of developments in the field have occurred necessitating improvements to this alignment tool.

The number and classification of true miRNAs has become controversial. miRBase, the central resource for miRNA curation, lists 2,588 human miRNAs in their latest version (v21), which has not been recently updated (9). Other manuscripts have listed thousands more putative novel miRNAs (10-12) including new passenger miRNA sequences of known miRNAs. However, the MirGeneDB group has indicated, using strict criteria, that only 858 human miRNAs exist, calling into question the continued search for novel miRNAs and perhaps the loose methods employed to designate short RNAs as miRNAs from deep RNA-seq data (13).

In recent years, there has also been an increased awareness and value placed on isomiRs. IsomiRs are categorized into three main classes: 5’ isomiRs, 3’ isomiRs and polymorphic isomiRs, with 5’ and 3’ isomiRs subclassified into templated and nontemplated modifications (14). The 5’ and 3’ isomiRs are the result of imprecise and alternative cleavage during the precursor miRNA (pre-miRNA) processing, post-transcriptional modifications, and/or editing by various post-transcriptional enzymes including exoribonucleases and nucleotidyl transferases (15-19). IsomiRs are beginning to be considered as more selective than just miRNA expression levels and must become well-characterized (20). True internal modifications (not technical artifacts) are generally the result of adenosine deaminase (ADAR) acting on RNA to cause an A to I modification (21) as noted in a variety of RNA species.

In response to these advancements, we now report major improvements in the 2.0 version of miRge. These include a highly-specific novel miRNA detector based on a machine learning algorithm and the ability to identify ADAR activity. Smaller revisions have been made to the algorithm to improve miRNA calling, increase flexibility of reporting and unification of the code base to Python for ease of programming and allowing for the implementation of miRge 2.0 into the bcbio-nextgen framework. We report the improvements and comparisons to other tools below.

## Materials and Methods

### Sequence databases and software dependencies

miRNA libraries were obtained from both miRBase.org (9) and MirGeneDB (13). mRNA and noncoding libraries were obtained from Ensembl (http://www.ensembl.org) unless otherwise noted. Human tRNAs were obtained from the Genomic tRNA Database (22). Human snoRNA was obtained from the snoRNABase (www-snorna.biotoul.fr). The redundant sequences in non-miRNA libraries were removed and the regions in these sequences which were identical to mature miRNAs were substituted with Ns. miRge 2.0 was written in Python (2.7.12) and utilizes the following tools and libraries: Bowtie (v1.1.1) (23), RNAfold (v2.3.5) (24), SAMtools (v1.5) (25), cutadapt (v1.11) (26), biopython (v1.68; http://biopython.org), sklearn (v0.18.1; http://scikit-learn.org), numPy (v1.11.0; http://www.numpy.org), SciPy (v0.17.0; https://www.scipy.org), pandas (v0.21.0; http://pandas.pydata.org), reportlab (v3.3.0; http://www.reportlab.com) and forgi (v0.20; https://viennarna.github.io/forgi). A installer incorporating all of these tools except Bowtie, SAMtools and RNAfold is included and the entire package is available through Bioconda. miRge 2.0 runs on a Linux platform (Ubuntu 16.04.3).

### miRge 2.0 Workflow

Figure 1 shows the workflow of miRge 2.0. In Figure 1, similar to the original miRge, the input FASTQ file(s) undergo prealignment steps of quality control, adaptor removal (cutadapt v1.11) and collapse into unique reads and their observed counts with subsequent merging across all unique samples (8). This file is then annotated against various search libraries, including mature miRNA, miRNA hairpin, mRNA, tRNA, snoRNA, rRNA, other non-coding RNA, and (optional) known RNA spike-in sequences (27,28). A full rationale of the method was given previously (8). The modifications of the search libraries are described in “Improvements of miRge 2.0” below. As an update from the original miRge approach, some alignment strategies were adjusted: 1) responding to a concern, only forward strand direction matching was allowed in the Bowtie step to search miRNAs with greater accuracy (29); 2) in the isomiR step, the Bowtie search was modified slightly from “bowtie -l 15 -5 1 -3 2 -n 2 –f” to “bowtie -5 1 -3 2 –v 2 –f –norc –best – S.”).

**Figure 1.**
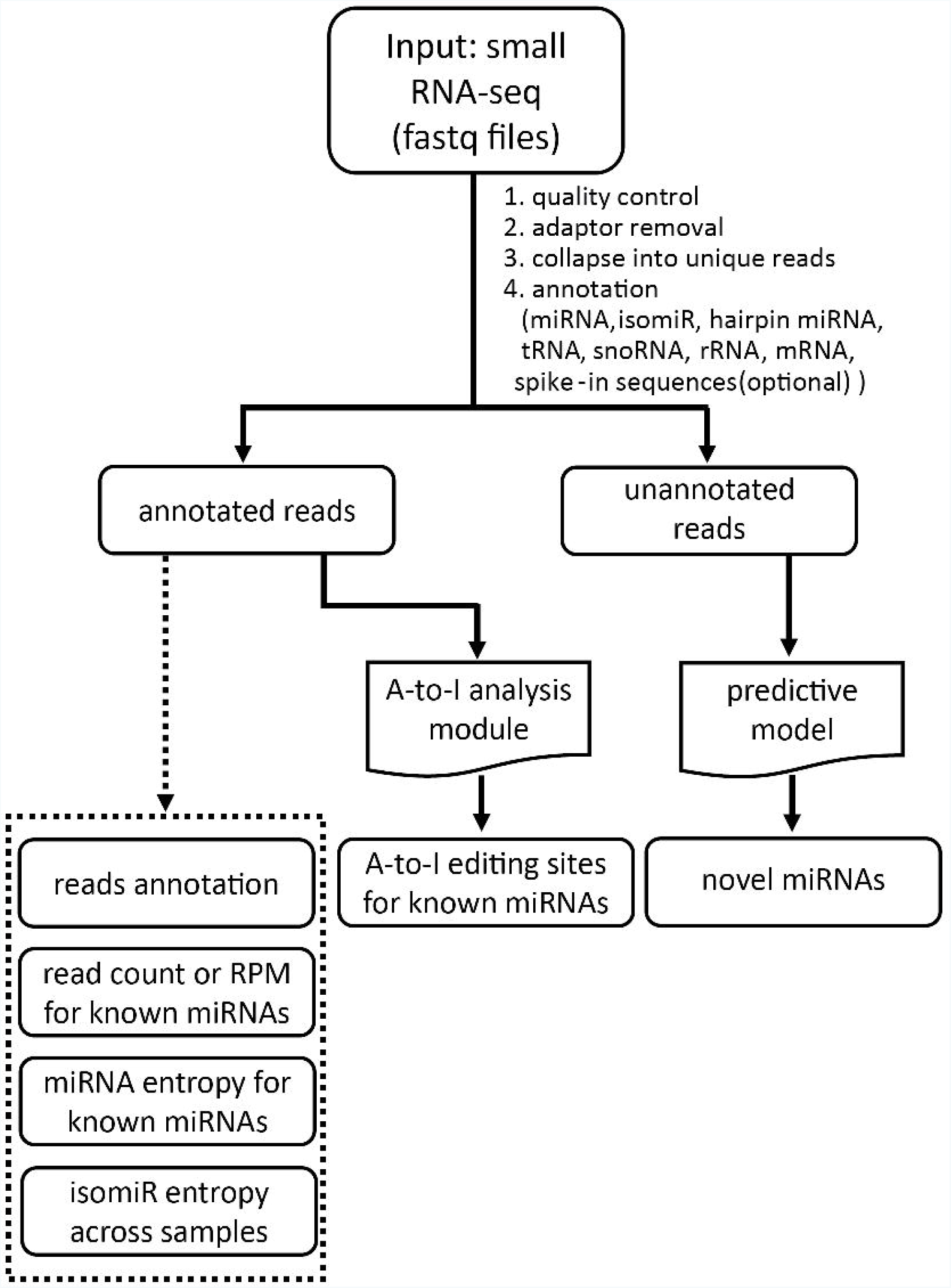
Workflow of miRge 2.0. It illustrates the flow chart from input to output. The modelsof A-to-I editing sites for known miRNAs and novel miRNAs detection are newly added functions, while the original outputs are shown in dashed box.

We addressed the effect of reads cross-mapping to more than one miRNA. We approach this by clustering the reads of the two or more similar miRNAs together (ex. hsa-miR-215-5p/192-5p). We made several improvements over the original miRge approach including systematically analyzing sequence similarity and merging miRNAs together if no mismatch is present in the main region of the miRNA. This was hand-curated and experimentally validated by repeated Bowtie alignments investigating random placement of reads.

Two new optional modules added in miRge 2.0 are the identification of ADAR A-to-I editing positions in the miRNAs and the search for putative novel miRNAs from unannotated reads (described below). Output files contain: 1) a .csv file containing all the annotated sequences; 2) two .csv files containing reads counts or reads per million (RPM) per miRNA; 3) an optional .csv file containing miRNA entropy and % canonical reads per miRNA; 4) an optional .csv file on the entropy of each isomiR across samples; 5) a .pdf report file containing an annotation log of the unique sequences identified across the entirety of the sample set analyzed along with per sample information on total reads, sequence length histograms, and the composition of the sample with respect to miRNA, mRNA, ncRNA, genomic, and unaligned reads; 6) an optional .gff file on the miRNAs and isomiRs (including CIGAR annotation) across samples; 7) an optional .csv file containing the identified significant A-to-I editing site in miRNAs and their proportion and adjusted p value; 8) an optional .pdf file showing a heat map of the A-to-I editing sites across samples; 9) an optional .csv report file of each sample containing the identified novel miRNAs; 10) Multiple .pdf files containing the structure of precursor miRNAs, the location, and reads alignment of novel miRNAs.

### Datasets to model novel miRNA detection

Sequencing datasets from 17 tissues in human and mouse (adrenal, bladder, blood, brain prefrontal cortex, colon, epididymis, heart, kidney, liver, lung, pancreas, placenta, retina, skeletal muscle, skin, testes and thyroid) were retrieved from Sequence Read Archive (Table 1). These samples were processed through miRge 2.0 to identify the different RNA species for machine learning controls. MirGeneDB miRNAs were used to assemble positive clusters (known miRNAs). RNAs in the categories of tRNA, snoRNA, rRNA or mRNA were used to assemble negative clusters (known non-miRNAs). Sequences in repeat elements were excluded. The details regarding the final selection of RNA species used are listed in “Generation of read clusters.” The collected miRNAs were further subselected by removing those that had less than 3 unique sequences, less than 10 overall reads, and are unable to form putative pre-miRNA structures. This yielded 12,048 and 7,795 known miRNAs (positive clusters) and 52,395 and 7,044 non-miRNAs (negative clusters) for the human and mouse datasets, respectively. To balance the positive and negative cluster data, 12,048 non-miRNA elements were randomly sampled from the original 52,395 in the human dataset and 7,044 miRNA elements were randomly sampled from the original 7,795 in the mouse dataset.

**Table 1.** Data sets for constructing the predictive model in human and mouse.

| Tissue Type | SRA References in human | SRA References in mouse |
| --- | --- | --- |
| Adrenal | SRR944031, SRR944034 | SRR3653309, SRR3653310 |
| Bladder | SRR333658, SRR333674 | SRR3652859, SRR3652860 |
| Blood | SRR837475, SRR837477 | SRR5241767, SRR5241768 |
| Brain Prefrontal Cortex | SRR1635903, ERR409900 | SRR3540303, SRR3540304 |
| Colon | SRR837839, SRR837842 | SRR1973865 |
| Epididymis | SRR384894 | NA |
| Heart | SRR553574, ERR038425 | SRR5832818, SRR5832819 |
| Kidney | SRR553575, ERR038420 | SRR3652244, SRR3652245 |
| Liver | ERR038413, ERR038410 | SRR5832837, SRR5832838 |
| Lung | SRR372648, SRR372650 | SRR5059366, SRR5059367 |
| Pancreas | ERR852097, ERR852099 | SRR1973869 |
| Placenta | SRR567637, SRR567638 | NA |
| Retina | ERR973611, ERR973613 | SRR1427160, SRR1427161 |
| Skeletal Muscle | SRR1635908, SRR1820680 | SRR3651659, SRR3651660 |
| Skin | SRR2174513, SRR2174517 | SRR3402126, SRR3402132 |
| Testes | SRR333680, SRR553576 | SRR1647951, SRR1647953 |
| Thyroid | SRR1291267, SRR1291269 | NA |

### Construction of the predictive model

Figure 2A illustrates the process of construction of the predictive model. 1) Annotated reads previously from a FASTQ file are classified into positive reads (miRNAs and isomiRs based on miRGeneDB) and negative reads (mRNAs and noncoding RNAs); 2) These raw sequence reads are mapped to the genome with perfect alignment and a new sequence cluster is generated based on their overlapping coordinates. The cluster sequences located at repeat regions are excluded. 3) All reads are then realigned to these putative mature miRNAs in an exact match round and single mismatch round using Bowtie. The most stable region of each cluster is extracted as a putative mature miRNA in the three steps, shown in Figure 2B. First all reads were aligned to the cluster; then we calculated the ratio of the read counts of each base along the cluster to the total read number for the cluster and finally set the start and end position of the putative miRNA as the first and last base with a ratio >0.8 of base position reads to total reads of the cluster. 4) We used these read structures to determine the optimal candidate pre-miRNA hairpins based on folding energy of the surrounding sequence. 5) The compositional features and the structural features of the pre-miRNAs were computed for each cluster. 6) A support vector machine (SVM) model, described below, was built to calculate the probability that a given candidate was a miRNA. 7) The probability of each putative mature miRNA is compared to the known positive or negative miRNA status of the read cluster to develop test statistics.

**Figure 2.**
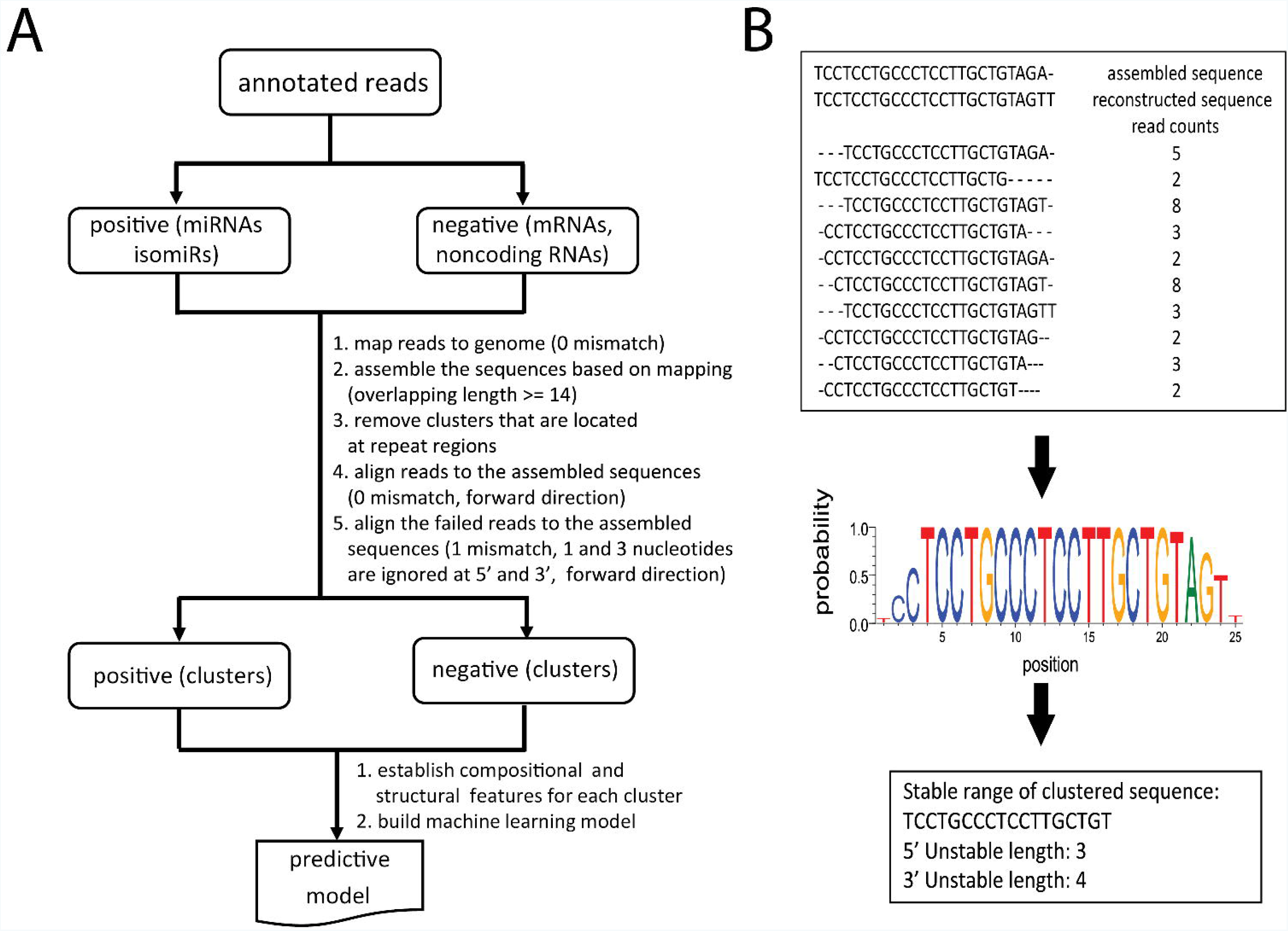
The process of construction of the predictive model. A) The building of the predictive model composed of data preparation, feature calculation, feature selection and machine learning model training. B) Schematic diagram of generating a stable range of clustered sequences in a cluster. The sequences in the cluster were aligned against the assembled sequence. The probability of the major nucleotide at each position was computed. A threshold of 0.8 was selected to determine the stable range of the cluster sequence.

### Generation of read clusters for annotated reads in model construction

After building the model with positive (known miRNAs) and negative (known non-miRNA) reads, we tested the model with reads that failed to map using our standard miRge alignments to known RNA libraries. From here, these unmapped reads were mapped to the human genome using Bowtie with 0 mismatches, seed length of 25 bp and alignment to 3 or fewer loci and then assembled based on coordinates. To form a cluster, two or more overlapping reads must have the same strand directionality with a minimum overlapped length of 14 bp. We removed assembled cluster sequences with length > 30 bp, a 6+ bp poly-A at the 3’ end, a 6+ bp poly-T at the 5’ end, or if they were located in a repetitive element region. All the reads were mapped to the assembled sequences with 0 mismatches, seed length of 25, and forward direction. The reads that did not align in this first step were mapped to the clusters in a less stringent manner, in which the first nucleotide and the last 3 nucleotides were ignored, up to 1 misaligned base pairs were allowed, seed length of 15 bp and forward strand direction was selected.

### Calculating compositional features of read clusters

miRNAs have a characteristic processing pattern to generate 5’ isomiRs and 3’ isomiRs. The 5’ end tends to stably begin at the same nucleotide, the 3’ end tends to be variable and nucleotide additions of uracils (U/T) or adenines (A) are frequently seen here. Non-miRNAs tend to not share these features, so this difference can be exploited. In order to codify these patterns, several features of read clusters were defined as follows: 1) the 5’ and 3’ unstable length of the cluster; 2) genome nucleotide proportion at the positions -3, -2, -1 of 5’ and +1, +2, +3, +4, +5, +6 of 3’ in the stable range of the cluster sequences; 3) A, T, C, and G percentages at the positions -3, -2, -1 of 5’ and +1, +2, +3, +4, +5, +6 of 3’ in the stable range of the cluster sequences. In addition, sequence type count, total read count and the proportion of reads that are an exact match to the cluster sequences were calculated as well.

### Generating precursor candidates for structural features

From the clustering process described above, genomic positions of the clusters (of known miRNAs known non-miRNAs, and unmapped reads) were obtained. The precursor (hairpin) candidate structures were generated as follows:

1. If a mapped cluster had no adjacent clusters, we determined a most-likely precursor (hairpin) structure. To do this, we generated two potential hairpin structures. We either added 20 nt upstream and 70 nt downstream or 70 nt upstream and 20 nt downstream of genomic sequence. Then, the secondary structure of the precursor was predicted by RNAfold. Any sequence without a hairpin secondary structure was removed at this step. After prediction, 5 nt was removed from the 20 nt side, and a matching length of sequence was removed from the 70 nt side, such that the hairpin had no overhang. We determined a 15 nt extended length is optimal for determining minimum free energy.
2. Precursor miRNA hairpin structures are discarded, if: a) a read cluster overlaps with the loop by more than 5 bp in the 5p-arm (no overlap is allowed on 3p-arm); b) the pruned pre-miRNA has no hairpin; c) the hairpin has less than 15 bindings in the total precursor structure; d) < 60% of nucleotides in the putative mature miRNA cluster are paired. If both precursor options remain, we chose the precursor with the lowest minimum free energy.
3. For those sequence clusters that had additional nearby clusters (within 44 bp), which could represent 5p and 3p arms, we approached these slightly differently. For these sequences, we assigned the neighbor state of each cluster sequence. To do this, we assigned the distance from the adjoining upstream sequence “seq1” to the target sequence “seq2” as D_1._ If there were three nearby sequence clusters, then we determined the distance from the adjoining downstream sequence “seq3” to the target sequence “seq2” as D_2_. If 9 <= D_1_<= 44 and the direction of “seq1” is equal to the direction of “seq2” or 9 <=D_2_<= 44 and the direction of “seq2” is equal to the direction of “seq3.” From each plausible scenario, we generated a precursor structure as described in 1) above and determined the optimal precursor based on the rules of 2) above.

### Prediction models for novel miRNA detection

We created a broad variety of features associated with cluster composition and precursor structures. All the features evaluated were listed in Supplemental Table 1. The discrimination power of each feature was ranked by its Minimum Redundancy Maximum Relevance (mRMR) score. We applied forward stepwise feature selection (30), to subselect the most informative features.

To test the model for robustness, the dataset was randomly split into training and test sets at the ratio of 4:1 in 10 replicates. The parameters of the estimator were optimized by 10 fold cross-validated grid-search over a parameter grid. The searching space of C and gamma in radial basis function kernel of SVM (31) were {0.0001, 0.001, 0.01, 0.1, 1.0, 10.0, 100.0, 1000}. The SVM model was implemented by scikit-learn Python package (http://scikit-learn.org). Matthews correlation coefficient (MCC) was used to evaluate the performance of the training model. The models were additionally tested on 12 rat samples (Supplemental Table 2).

### A-to-I editing analysis

We utilized the mapped output file to identify all reads corresponding to each miRNA for A-to-I editing, as noted as an A to G change. First, the reads were aligned against the genome with the last two nucleotides at the 3’ end trimmed and allowance of up to one mismatch. Here, we demanded unique best hits (i.e. a read that can’t be aligned to other locations in the genome with the same number of mismatches). Then, for the retained reads that belong to one miRNA and its isomiR, all nucleotide positions in the canonical miRNA, except the terminal 5 bp were screened for A to G changes based on a binomial test considering the expected sequencing error rate (0.1%), as described (32). A Benjamini-Hochberg-corrected P-value (33) was calculated for each site on the miRNA. The A-to-I editing level was defined as the proportion of the mapped reads containing the edited nucleotide relative to the total mapped reads at the given location. Finally, we excluded the putative A-to-I signals if: 1) the locations where similar miRNA families or miRNA SNPs that have A/G differences could be mistaken for A-to-I changes (i.e. nucleotide position 19 in let-7a-5p and let-7c-5p which differ only by an A/G variation and miR- 548al which has a SNP (A-to-G) at position 8 with the frequency of 0.18); 2) the 455 miRNAs found in repeat elements which could give false positives (i.e. miR-6503-3p is located in a MTL1D long terminal repeat.); 3) the miRNAs where the RPM of the canonical sequence is less than 1; 4) the miRNAs where the corresponding one nucleotide switched sequence (A to G) can be aligned to more than 1 location in the genome with trimming the last two nucleotides at 3'.

### Comparison to other novel miRNA tools

Currently, miRDeep2 (34) and miRAnalyzer (35) are two prevailing tools for the prediction of novel miRNAs. In our annotation comparison study, default parameters were utilized except that the ‘-l’ was set to be 17 in the mapper.pl for miRDeep2 and default parameters were utilized in miRAnalyzer. In our prediction comparison study, new FASTQ files were generated from the unmapped read data of an original miRge run. Default parameters were utilized when running miRDeep2 and miRAnalyzer. Two metrics of novel miRNAs were used to compare three tools: PhyloP score and quality score. Basewise conservation scores across miRNAs were calculated from PhyloP data downloaded from http://hgdownload.cse.ucsc.edu/goldenPath/hg38/phyloP20way/ (36) using the PHAST package (37). For each miRNA, the mean of PhyloP values across its length was calculated. The quality scores for each miRNA by each tool was defined by: 1 - (ranking percentile by the tool).

### Hardware

All processing was performed on a workstation with 56 CPUs (dual Intel(R) Xeon(R) E5-2690 v4 at 2.60GHz) and 256GB DDR4-RAM. Novel miRNA modelling was performed using 32 CPUs. For speed testing, the number of CPUs in running original miRge, miRge 2.0, miRDeep2 and miRAnalyzer were 5, 5, 1 and 1, respectively. Due to a java incompatibility on the workstation, miRAnalyzer was run on a desktop with 4 CPUs (Intel(R) Core(TM) i7-6700 CPU at 3.40GHz) and 16GB DDR4-RAM.

## Results

### Improvements of miRge 2.0

The major improvements of miRge 2.0 consist of a novel miRNA detection method, improved alignment parameters, and the reporting of A-to-I changes in the sequence. These are described below, while smaller improvements are reported here. By changing the search parameters, miRge 2.0 is able to annotate reads more precisely. In human data, using the miRBase library, we search for 2,741 miRNAs of which 134 are merged due to a similarity of their sequences. miRge 2.0 provides an optional GFF file report, which includes isomiR CIGAR values. These can be used for isomiR-driven analyses. Additionally, the GFF data file is easily incorporated into other analysis pathway software including the bcbio-nextgen framework. miRge 2.0 also generates a .csv and .pdf file report of summary statistics; replacing a html report which was more difficult to process for tabular information. We also made several revisions to the search libraries. For the miRBase-based alignment search, we included additional SNP information in the miRNA library based on the updated miRNASNP database (38). We have also included 153 5p or 3p miRNAs that are the complement of known miRBase miRNAs for which the passenger strand was detected recently (11). Thus we have expanded our miRBase search library from 2,588 miRNAs in our original method to 2,741 miRNAs currently. We have also built a MirGeneDB-based alignment library that is corrected for SNP information for those investigators seeking this more specific set of miRNAs. For any alignment, we have added an optional spike-in RNA library search based on two popular sources of spike-in normalization (27,28). This search can easily be expanded to capture newer spike-in normalization methods as they appear. All options to call in miRge 2.0 are shown in Supplemental Table 3.

### Speed and annotation comparison of original miRge and miRge 2.0

We performed tests of speed and annotation function of miRge 2.0 using six datasets. Both miRBase and MirGeneDB based libraries were analyzed although novel miRNA detection and A-to-I analysis were not performed. We found miRge 2.0 required 10%-20% more processing time than the original miRge (totaling approximately 1-2 minutes more time) due to increased and larger bowtie searching libraries and adjusted searching parameters (Table 2). The number of detected miRNAs was slightly decreased as well. The alignment speed was essentially the same as miRAnalyzer and significantly faster than miRDeep2. The discovery of novel miRNAs is more time and memory intensive, as expected. For the dataset SRR553572 with 25.7 million reads, to identify novel miRNAs, the calculation time and maximum memory consumption were 1.2 h and 6.7 GB RAM respectively.

**Table 2.**
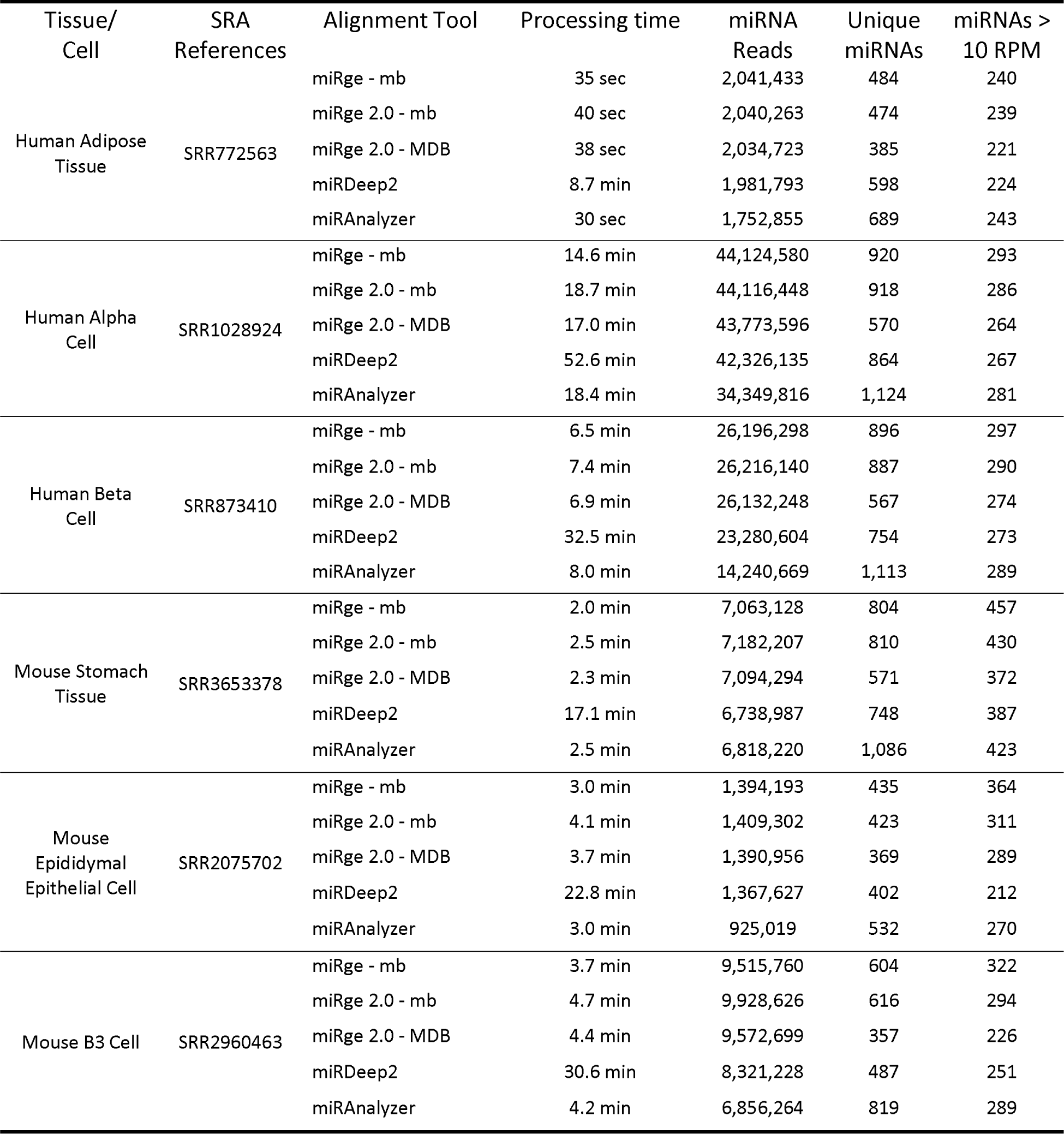
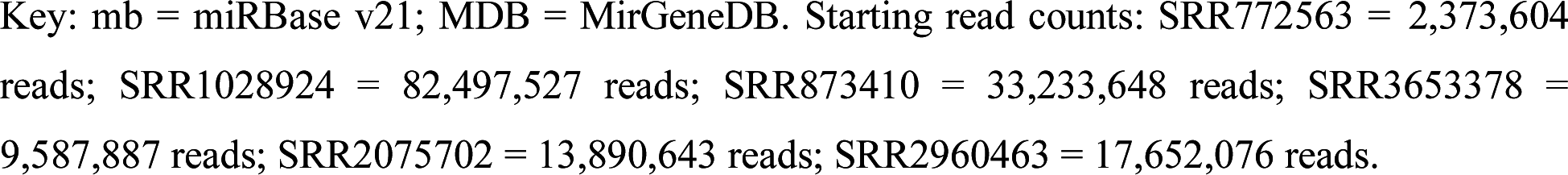
Data sets for constructing the predictive model in human and mouse.

| Tissue/<br>Cell | SRA<br>References | Alignment Tool | Processing time | miRNA<br>Reads | Unique<br>miRNAs | miRNAs ><br>10 RPM |
| --- | --- | --- | --- | --- | --- | --- |
| Human Adipose<br>Tissue | SRR772563 | miRge - mb | 35 sec | 2,041,433 | 484 | 240 |
|  |  | miRge 2.0 - mb | 40 sec | 2,040,263 | 474 | 239 |
|  |  | miRge 2.0 - MDB | 38 sec | 2,034,723 | 385 | 221 |
|  |  | miRDeep2 | 8.7 min | 1,981,793 | 598 | 224 |
|  |  | miRAnalyzer | 30 sec | 1,752,855 | 689 | 243 |
| Human Alpha<br>Cell | SRR1028924 | miRge - mb | 14.6 min | 44,124,580 | 920 | 293 |
|  |  | miRge 2.0 - mb | 18.7 min | 44,116,448 | 918 | 286 |
|  |  | miRge 2.0 - MDB | 17.0 min | 43,773,596 | 570 | 264 |
|  |  | miRDeep2 | 52.6 min | 42,326,135 | 864 | 267 |
|  |  | miRAnalyzer | 18.4 min | 34,349,816 | 1,124 | 281 |
| Human Beta<br>Cell | SRR873410 | miRge - mb | 6.5 min | 26,196,298 | 896 | 297 |
|  |  | miRge 2.0 - mb | 7.4 min | 26,216,140 | 887 | 290 |
|  |  | miRge 2.0 - MDB | 6.9 min | 26,132,248 | 567 | 274 |
|  |  | miRDeep2 | 32.5 min | 23,280,604 | 754 | 273 |
|  |  | miRAnalyzer | 8.0 min | 14,240,669 | 1,113 | 289 |
| Mouse Stomach<br>Tissue | SRR3653378 | miRge - mb | 2.0 min | 7,063,128 | 804 | 457 |
|  |  | miRge 2.0 - mb | 2.5 min | 7,182,207 | 810 | 430 |
|  |  | miRge 2.0 - MDB | 2.3 min | 7,094,294 | 571 | 372 |
|  |  | miRDeep2 | 17.1 min | 6,738,987 | 748 | 387 |
|  |  | miRAnalyzer | 2.5 min | 6,818,220 | 1,086 | 423 |
| Mouse<br>Epididymal<br>Epithelial Cell | SRR2075702 | miRge - mb | 3.0 min | 1,394,193 | 435 | 364 |
|  |  | miRge 2.0 - mb | 4.1 min | 1,409,302 | 423 | 311 |
|  |  | miRge 2.0 - MDB | 3.7 min | 1,390,956 | 369 | 289 |
|  |  | miRDeep2 | 22.8 min | 1,367,627 | 402 | 212 |
|  |  | miRAnalyzer | 3.0 min | 925,019 | 532 | 270 |
| Mouse B3 Cell | SRR2960463 | miRge - mb | 3.7 min | 9,515,760 | 604 | 322 |
|  |  | miRge 2.0 - mb | 4.7 min | 9,928,626 | 616 | 294 |
|  |  | miRge 2.0 - MDB | 4.4 min | 9,572,699 | 357 | 226 |
|  |  | miRDeep2 | 30.6 min | 8,321,228 | 487 | 251 |
|  |  | miRAnalyzer | 4.2 min | 6,856,264 | 819 | 289 |

Key: mb = miRBase v21; MDB = MirGeneDB. Starting read counts: SRR772563 = 2,373,604 reads; SRR1028924 = 82,497,527 reads; SRR873410 = 33,233,648 reads; SRR3653378 = 9,587,887 reads; SRR2075702 = 13,890,643 reads; SRR2960463 = 17,652,076 reads.

### A-to-I editing analysis

To evaluate the accuracy of A-to-I editing analysis, we performed A-to-I analysis using a pooled human brain sample (SRR095854) and compared the results to prior published data on this sample (32). We identified 19 significant A-to-I modification sites compared to 16 reported in the reference paper. Comparing the two sets of results, the adjusted R^2^ of A-to-I proportion of these shared 16 sites was 0.96 and the slope of the linear regression was 0.99 indicating high reproducibility between our method and the established method (Figure 3A). We then performed a new A-to-I editing analysis across colon tissue (Sequence Read Archive samples: SRR837842 and SRR837839), colon epithelial cells (SRR5127219), colon cancer (SRR1646473 and SRR1646493), and the colon cancer cell lines DKO1 (SRR1917324), DLD1 (SRR1917336) and DKS8 (SRR1917329). Significant miRNA editing sites with A-to-I percentage ≥ 1% in at least one sample were shown in Figure 3B, with the data indicating differences between tumor and normal cells in ADAR activity (39).

**Figure 3.**
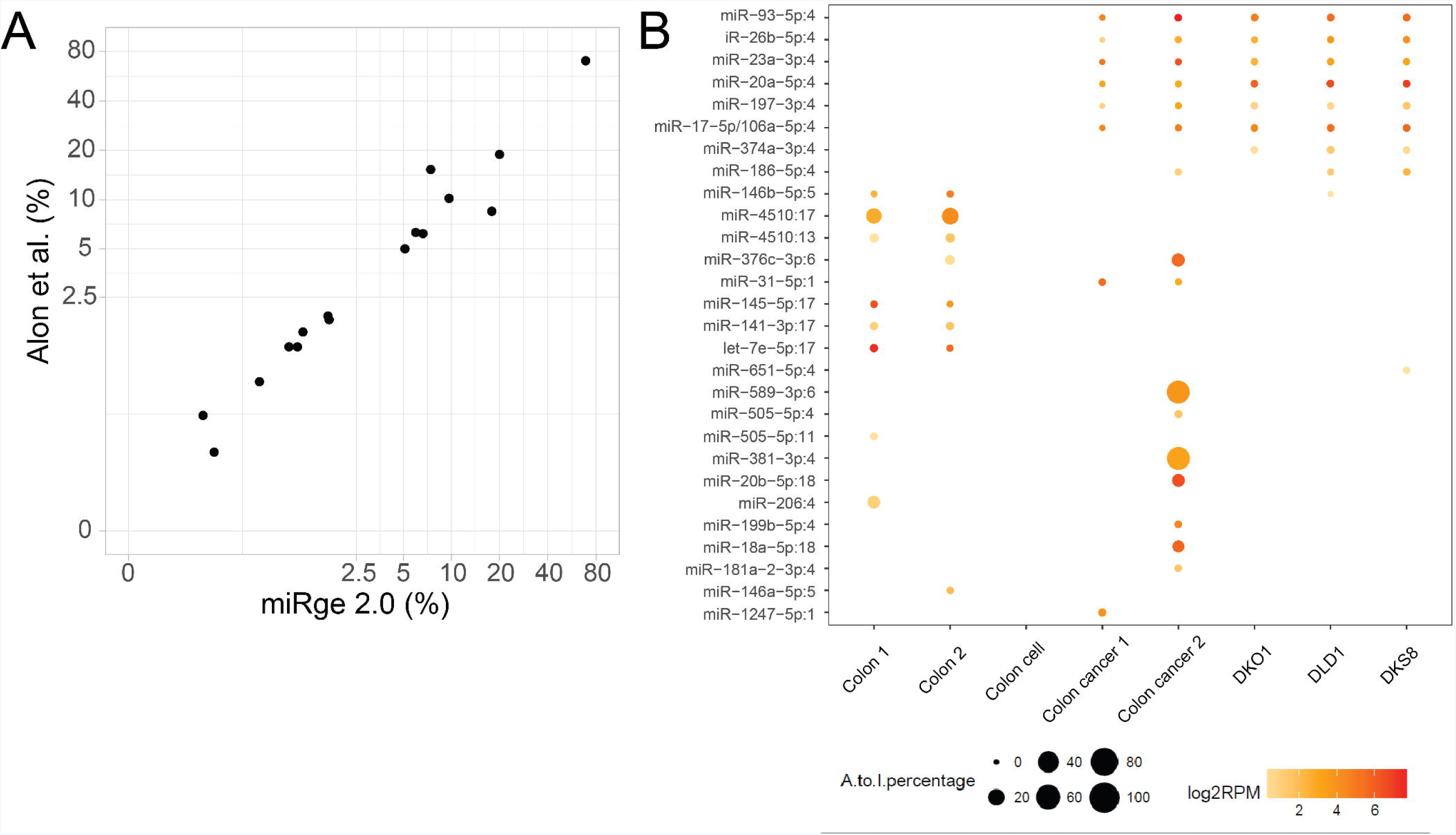
A-to-I analysis. A) The A-to-I proportion of the sites is strongly correlated with a reference dataset analysis with adjusted R^2^ of 0.96 in the log-log plot. B) The output of miRge 2.0 showing an illustrated heat map of miRNA A-to-I editing sites across colon tissue, primary colon cell, colon cancer tissue and colon cancer cells from multiple sources.

### Validation of the predictive model

To determine the optimal number of features to use in the human and mouse predictive model, the MCC for the training and test sets for the top 40 ranked features based on mRMR scores are shown in Figure 4. For human data, when the number of features reached 21, the mean value of MCC of training and test set approached the maximum and became stable. These top features are listed in Table 3. Among them, there are 11 precursor miRNA structural features and 10 compositional features. The ultimate model was constructed using these selected features. We used 32 human cell data sets to validate the model as an external test set. The positive and negative miRNAs were generated through the same process described above. The predictive result is shown in Table 4. The mean of MCC is 0.94, indicating that the performance of the model in the external test set is good.

**Figure 4.**
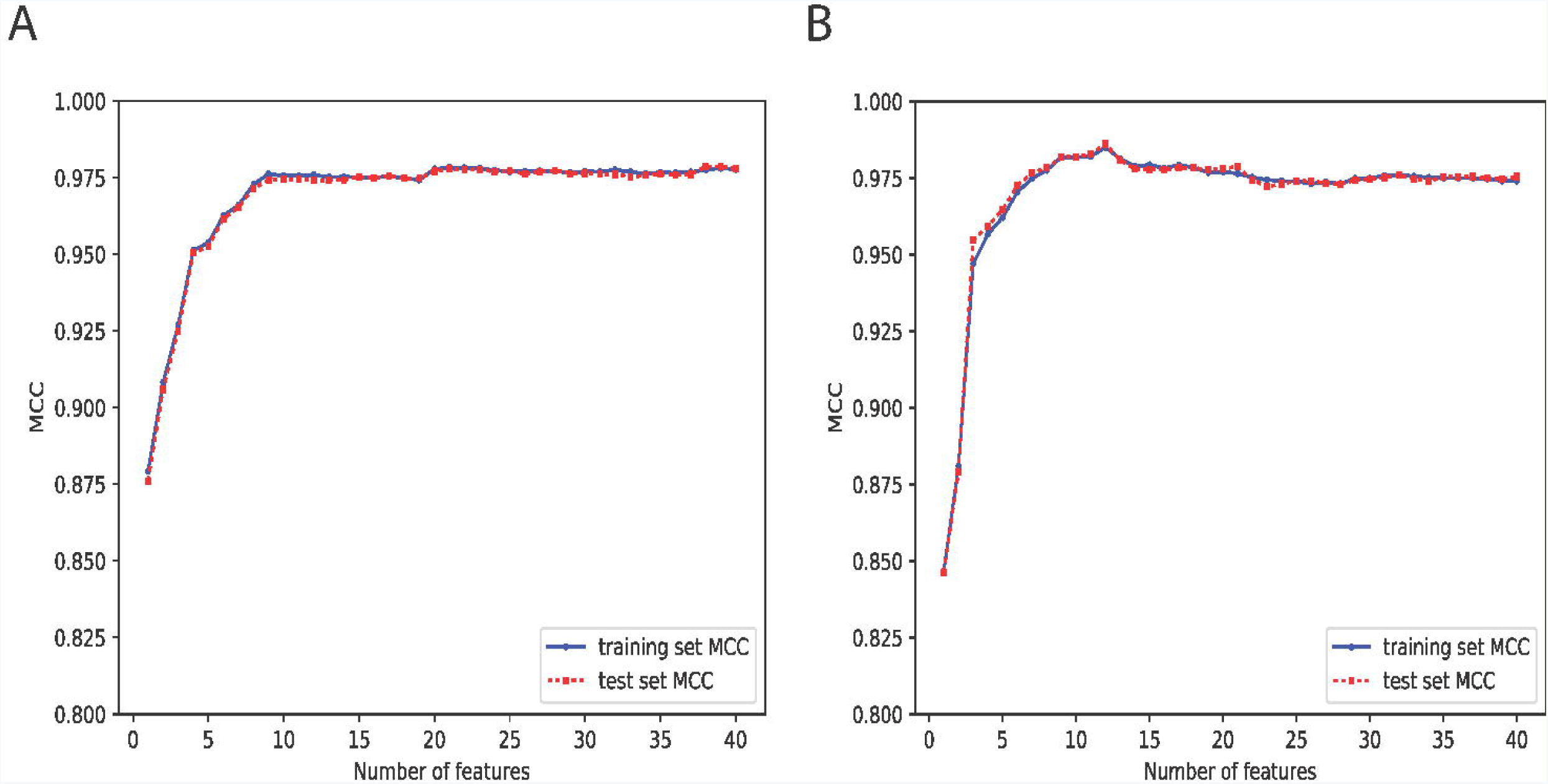
Model performance on top 40 features for training and test sets for human (Figure 4A)and mouse (Figure 4B) miRNA discovery. Each dot stands for the mean value of Matthews correlation coefficient (MCC).

**Table 3.**
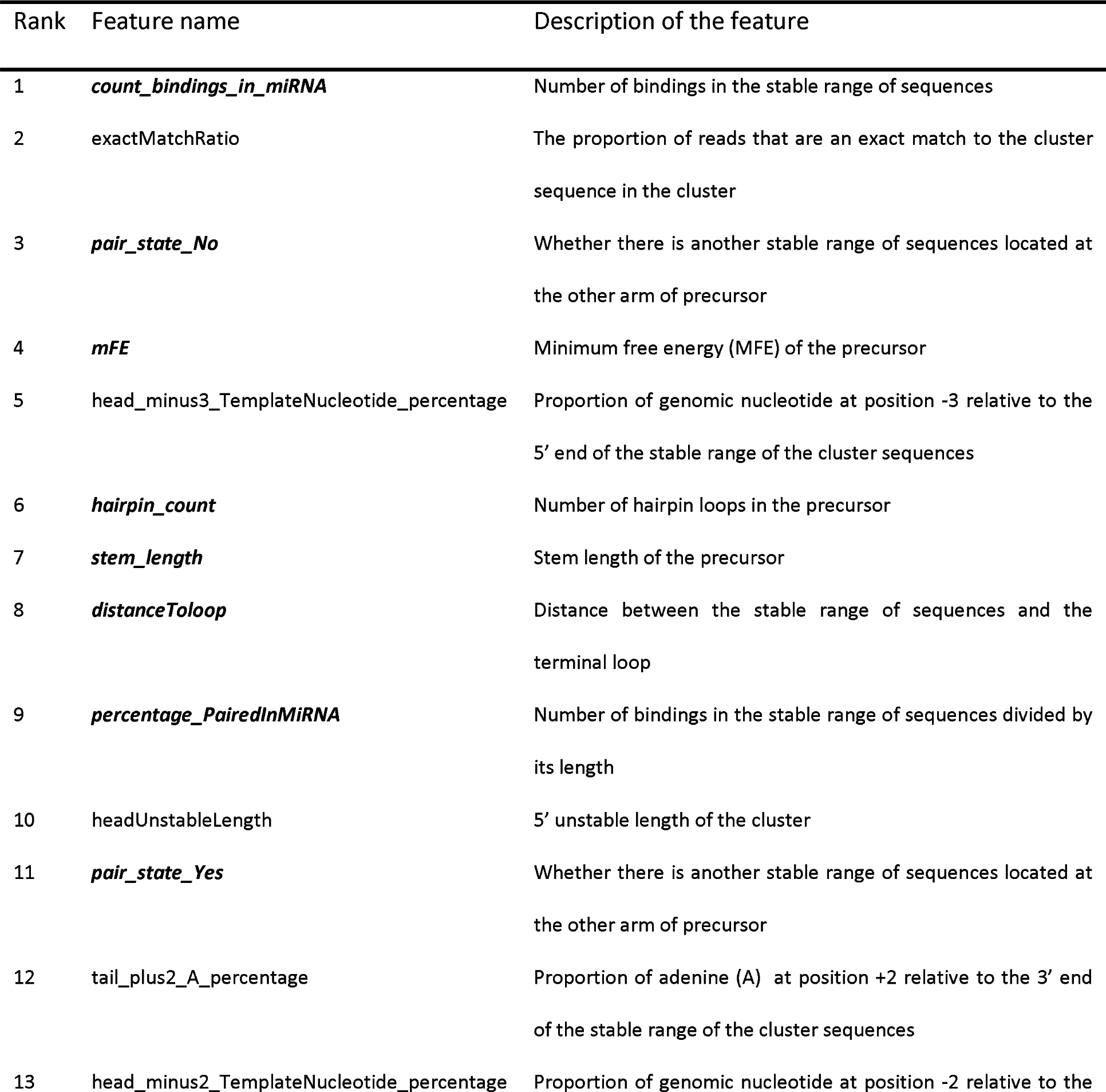
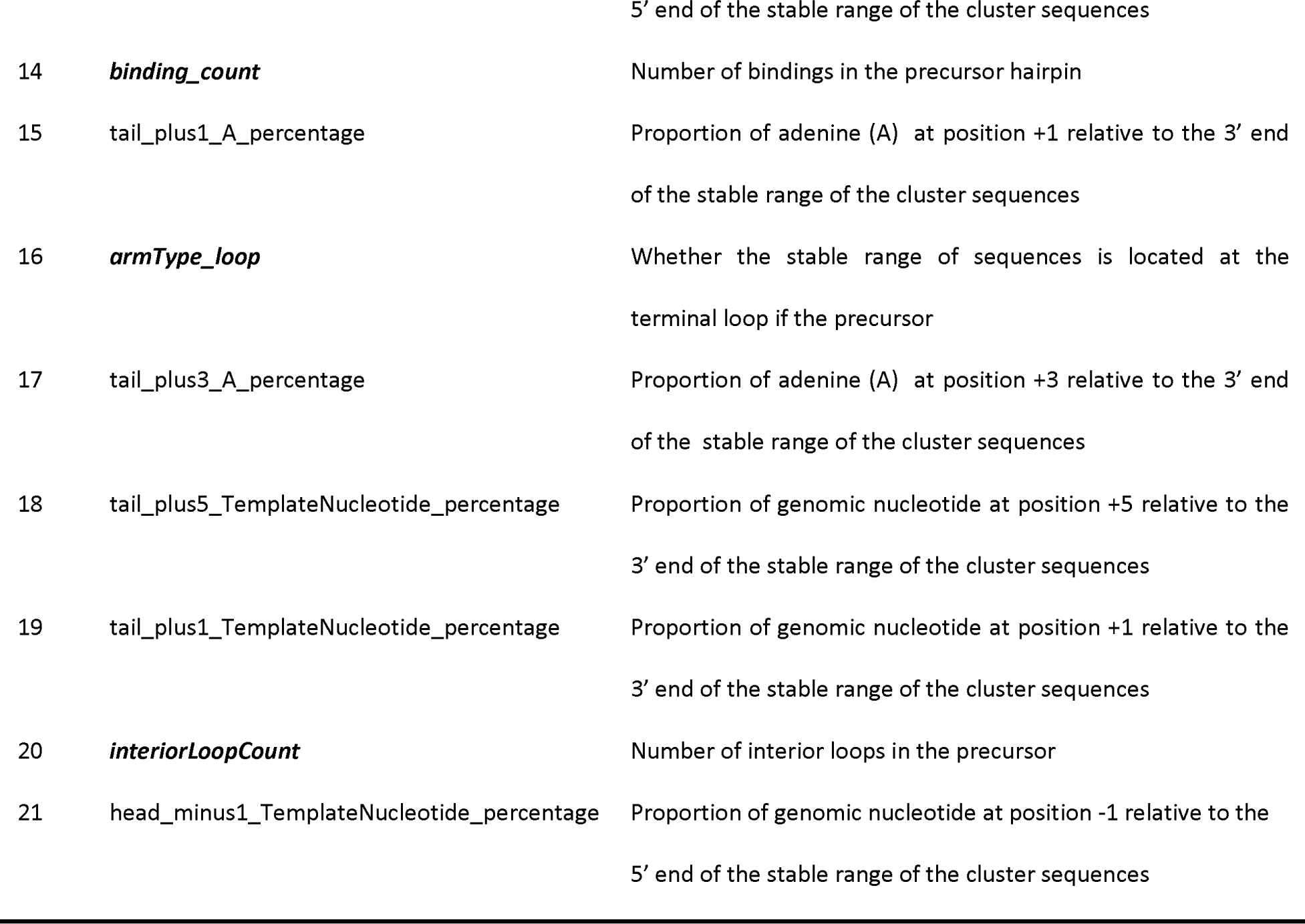
Top 21 features in human predictive model. Hairpin structural features are labeled in bold/italics, while read compositional features are not.

**Table 4.** Predictive results of 32 human cell data as external test sets by human model.

| Cell Type | SRA References | AUC | Precision | Recall | MCC |
| --- | --- | --- | --- | --- | --- |
| Fibroblast Aorta Adventitia | SRR5127206 | 0.995 | 0.983 | 0.963 | 0.945 |
| Smooth Muscle Cell Aorta | SRR5127217 | 0.994 | 0.981 | 0.961 | 0.938 |
| Astrocyte | SRR5127214 | 0.994 | 0.98 | 0.968 | 0.949 |
| Smooth Muscle Cell Bladder | SRR5127215 | 0.992 | 0.971 | 0.963 | 0.936 |
| Fibroblast Dermal (Adult) | SRR5127205 | 0.995 | 0.983 | 0.974 | 0.95 |
| Fibroblast Dermal (Neonatal) | SRR5127225 | 0.995 | 0.989 | 0.96 | 0.942 |
| Epithelium Keratinocyte (Adult) | SRR5127203 | 0.994 | 0.977 | 0.962 | 0.934 |
| Epithelium Keratinocyte (Neonatal) | SRR5127208 | 0.993 | 0.975 | 0.942 | 0.923 |
| Endothelial Aortic | SRR5139121 | 0.988 | 0.975 | 0.932 | 0.915 |
| Endothelial Umbilical vein | SRR5127213 | 0.993 | 0.981 | 0.954 | 0.926 |
| Epithelium Bronchial | SRR5127216 | 0.988 | 0.974 | 0.951 | 0.935 |
| Chondrocyte | SRR5127229 | 0.995 | 0.985 | 0.959 | 0.944 |
| Endothelial Microvascular | SRR5127201 | 0.991 | 0.973 | 0.957 | 0.945 |
| Fibroblast Cardiac | SRR5127236 | 0.992 | 0.983 | 0.945 | 0.94 |
| Melanocyte | SRR5127207 | 0.995 | 0.99 | 0.981 | 0.954 |
| Epithelium Mammary | SRR5127224 | 0.99 | 0.976 | 0.941 | 0.927 |
| Epithelium Prostate | SRR5127212 | 0.992 | 0.975 | 0.961 | 0.948 |
| Epithelium Renal Cortex | SRR5127204 | 0.988 | 0.966 | 0.948 | 0.927 |
| Epithelium Renal Proximal | SRR5127230 | 0.992 | 0.978 | 0.949 | 0.936 |
| Stromal cell Prostate | SRR5127226 | 0.991 | 0.976 | 0.963 | 0.94 |
| Myoblast Skeletal Muscle | SRR5127218 | 0.99 | 0.974 | 0.956 | 0.932 |
| Epithelium Intestinal | SRR5127223 | 0.994 | 0.985 | 0.973 | 0.957 |
| Myofibroblast | SRR5127220 | 0.991 | 0.987 | 0.965 | 0.95 |
| Smooth Muscle Cell Prostate | SRR5127222 | 0.991 | 0.978 | 0.961 | 0.943 |
| Neuron Dopaminergic | SRR5127234 | 0.982 | 0.963 | 0.922 | 0.916 |
| Neuron Cortical | SRR5127209 | 0.986 | 0.968 | 0.917 | 0.917 |
| Mesangial | SRR5127221 | 0.996 | 0.986 | 0.971 | 0.948 |
| Osteoblast | SRR5127233 | 0.997 | 0.986 | 0.955 | 0.946 |
| Fibroblast Periodontal ligament | SRR5127227 | 0.994 | 0.986 | 0.962 | 0.946 |
| Epithelium Renal | SRR5127235 | 0.992 | 0.989 | 0.954 | 0.936 |
| Epithelium Retinal Pigment | SRR5127210 | 0.994 | 0.988 | 0.974 | 0.959 |
| Skeletal Muscle Cell | SRR5127202 | 0.995 | 0.984 | 0.96 | 0.936 |
| Mean |  | 0.992 | 0.98 | 0.956 | 0.939 |
| Std dev |  | 0.003 | 0.007 | 0.014 | 0.012 |

Meanwhile, in the mouse predictive model, the optimal number of features are 12 which is shown in Supplemental Table 4. These 12 features are a subset of the 21 human features used. The performance of mouse model towards 19 mouse cell datasets are shown in Supplemental Table 5 where the mean of MCC is 0.93, indicating that the mouse model performed well on the external dataset.

### Comparison with other novel miRNA detection tools

Using miRge 2.0, we identified 302 RNA species that are putative novel miRNAs from 32 cell types (11). Referring to these sequences as novel miRNAs, without further validation, may be incorrect terminology. However, without other terminology for these small “true miRNAs” or “miRNA-like RNA species,” we will refer to them as putative novel miRNAs. We then used the same unmapped reads generated from miRge 2.0 as input for miRDeep2 and miRAnalyzer. They predicted 1,975 and 18,168 putative novel miRNAs respectively. After thresholding the data from those two tools to the same parameters as miRge 2.0 (≥10 total reads, ≥3 sequences, etc.), there were 312 and 391 putative novel miRNAs remaining. As shown in Figure 5A, a Venn diagram depicts the overlap among miRge 2.0, miRDeep2 and miRAnalyzer, showing 129 novel miRNAs shared between the three methods. We then calculated the mean PhyloP scores as a measure of nucleotide conservation across primates for the novel miRNAs (Figure 5B). More conservation was noted for the shared novel miRNAs (0.14) compared to miRAnalyzer (0.013) and miRDeep2 (-0.036). Conservation was equivocal between the shared novel miRNAs and the miRge 2.0 novel miRNAs (0.15) As all three tools give a quality score to each novel prediction, we compared these values for miRNAs found shared vs. those unique to each method. As shown in Figure 5C, the overlapped miRNAs ranked higher in quality for each method, further suggesting these 129 are the optimal putative novel miRNAs from the group. The full list of putative novel miRNAs generated by the three tools are available in Supplemental Table 6.

**Figure 5.**
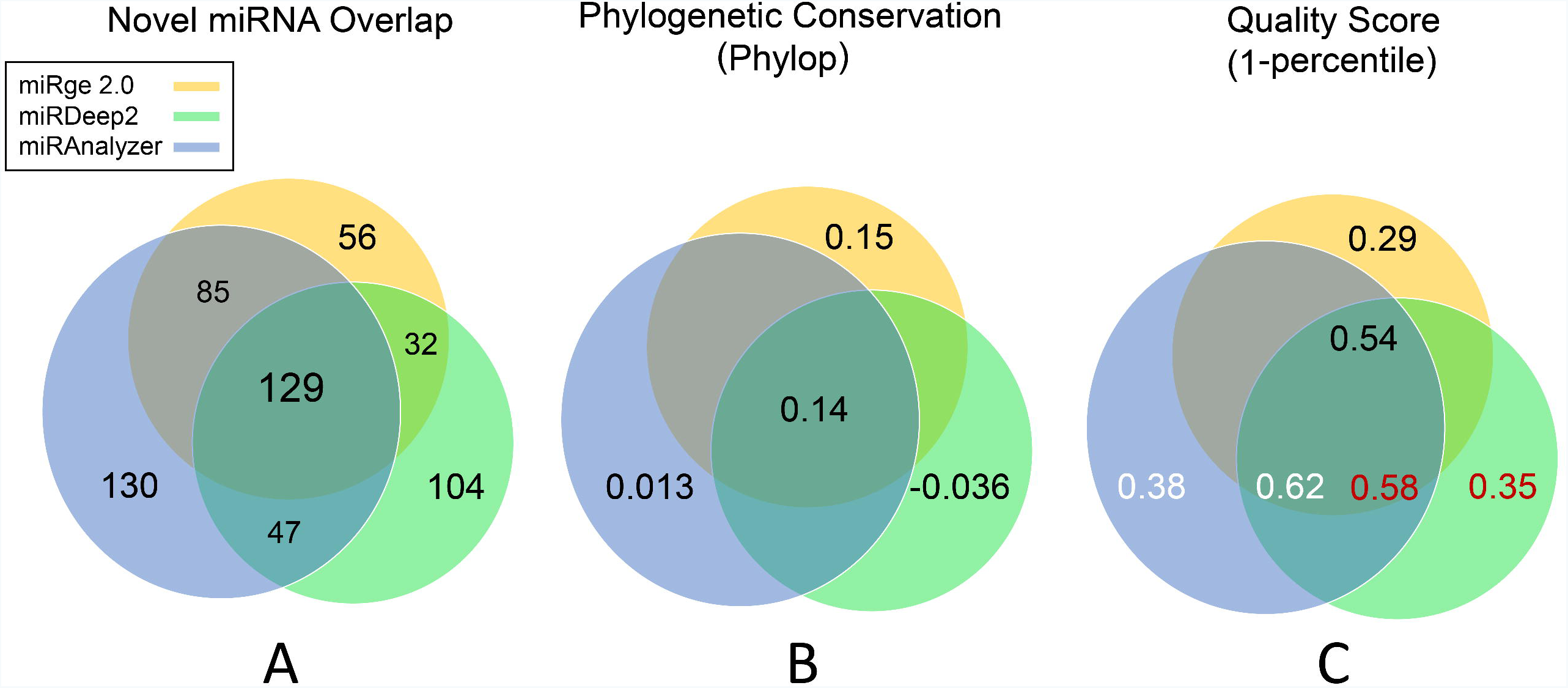
Venn diagram for novel miRNAs predicted by miRge 2.0, miRDeep2, and miRAnalyzer. A) Overlapped novel miRNAs among the three tools. B) The average basewise conservation scores across novel miRNAs. C) The average Quality score across novel miRNAs among the three tools.

### Comparison between the human model and mouse model

We used our SVM model to create an optimal novel miRNA tool for both human and mouse. We questioned how well those tools could predict novel miRNAs in other species. We utilized both the human and mouse models on the 12 rat miRNA samples shown in Supplemental Table 2. Using known miRNAs and known non-miRNAs, we found the average MCC for the rat samples to be equivalent when using the human model (0.96) than when using the mouse model (0.95). Therefore, either model seems useful for finding novel miRNAs in at least other mammals.

## Discussion

In light of the positive and negative feedback we received for our original miRge tool, we set about to make an improved 2.0 version. miRge 2.0 has a more robust search, better overall output reporting, more run options, and new parameters for novel miRNA detection and A-to-I editing detection. It can be implemented within the bcbio-nextgen framework to better integrate with other software tools. It still remains one of the fastest options for alignment and can multiplex multiple samples in a single run. The new novel miRNA detection tool has reasonable requirements for RAM and can be used widely.

Our data suggests the miRge 2.0 novel miRNA detection tool is more robust than the earlier tools miRDeep2 and miRAnalyzer. We believe that our unique use of compositional features improved miRNA discovery. The strategy of removing clusters if there were > 3 unique sequences, > 10 overall reads, and/or lacked a pre-miRNAs structure also reduced the false positive rate. We caution though, that these are putative novel miRNAs and should not be thought of as bona fide miRNAs unless they meet additional parameters (13). We are also wondering if a novel detection tool built for one mammalian species could be used to detect putative novel miRNAs in other species. Our human and mouse models assayed with the rat data indicates, that, indeed, at least among mammalia, our tool is robust.

We have also tried to make miRge 2.0 more robust to current concerns of the community. Many authors have argued that miRBase—the online repository for miRNAs —is riddled with false positive miRNAs (40-42). Therefore, we have built a MirGeneDB-based alignment library, incorporating SNPs, for 6 species to cater to those investigators seeking a better-defined set of miRNAs. We have reported concerns with using reads per million miRNA reads (RPM) as a normalization tool (43). Therefore, we have added an optional spike-in RNA library search step for spike-in normalization. Spike-in for miRNA RNA-seq is still in its infancy, so this step can easily be expanded/modified to account for newer spike-in normalization methods. Currently, the sequence libraries of human, mouse, rat, nematode, fruitfly and zebrafish datasets are provided, but miRge 2.0 can be used by individual users to investigate any species by constructing the sequence libraries to incorporate in the miRge 2.0 workflow using our miRge_bowtie_build.py tool.

In our original miRge tool, we accepted that reads could randomly align to highly similar miRNAs, e.g. miR-192-5p and miR-215-5p; thus we reported those together as miR-215-5p/192-5p reads. The cross-mapping of sequencing reads can create false alignments that may be interpreted as sequence or expression alterations which can occur in other alignment tools, as other tools have generally not hand-curated their alignment libraries. Our improvements in miRge 2.0 optimize the number of miRNAs that are clustered together to reduce these random alignment challenges.

With the interest in ADAR activity and A-to-I changes in RNAs, we have added a feature to miRge 2.0 to capture this information. miRge 2.0 performs robustly in identifying these ADAR sites, comparable to other stand-alone programs.

In summary, miRge 2.0 is an update of our original miRNA alignment tool that more comprehensively and more robustly analyzes miRNA sequencing data. We believe the improvements in miRge 2.0 will be useful to a wide range of scientists who are interested in interpreting small RNA-seq data for miRNA expression patterns.

## Acknowledgements

The authors thank Lorena Patano and Arun H. Patil for assistance in troubleshooting program installation.

